# Maximum intrinsic rate of population increase in sharks, rays, and chimaeras: the importance of survival to maturity

**DOI:** 10.1101/051482

**Authors:** Sebastián A. Pardo, Holly K. Kindsvater, John D. Reynolds, Nicholas K. Dulvy

**Affiliations:** Earth to Ocean Research Group, Department of Biological Sciences, Simon Fraser University, Burnaby, BC, Canada; Department of Ecology, Evolution, and Natural Resources, Rutgers University, New Brunswick, NJ, USA

**Keywords:** elasmobranch, extinction risk, demography, data-poor, population growth rate, recovery potential

## Abstract

The maximum intrinsic rate of population increase *r*_*max*_ is a commonly estimated demographic parameter used in assessments of extinction risk. In teleosts, *r*_*max*_ can be calculated using an estimate of spawners per spawner, but for chondrichthyans, most studies have used annual reproductive output *b* instead. This is problematic as it effectively assumes all juveniles survive to maturity. Here, we propose an updated *r*_*max*_ equation that uses a simple mortality estimator which also accounts for survival to maturity: the reciprocal of average lifespan. For 94 chondrichthyans, we now estimate that *r*_*max*_ values are on average 10% lower than previously published. Our updated *r*_*max*_ estimates are lower than previously published for species that mature later relative to maximum age and those with high annual fecundity. The most extreme discrepancies in *r*_*max*_ values occur in species with low age at maturity and low annual reproductive output. Our results indicate that chondrichthyans that mature relatively later in life, and to a lesser extent those that are highly fecund, are less resilient to fishing than previously thought.

## 1 Introduction

The rate of increase is a fundamental property of populations that arises from birth and death rates. A commonly used metric for guiding assessments of extinction risk and setting limit reference points is the maximum intrinsic rate of population increase *r*_*max*_; it reflects the productivity of depleted populations where density-dependent regulation is absent (Myers and Mertz, 1998; Myers et al., 1997). When population trajectories are lacking, *r*_*max*_ is useful for evaluating a species’ relative risk of overexploitation (Dulvy et al., 2014) as it is equivalent to the fishing mortality that will drive a species to extinction, *F*_*ext*_ (Myers and Mertz, 1998). A fundamental parameter in calculating *r*_*max*_ is the product of survival to maturity *l*_*αmat*_ and annual fecundity *b*. Fisheries biologists studying teleost fishes often calculate it based on lifetime spawners per spawner (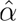), which is related to the slope near the origin of a stock-recruitment relationship (Denney et al., 2002; Dulvy et al., 2004; Hutchings et al., 2012). In other words, the spawners per spawner incorporates juvenile survival and approximates *l*_*αmat*_*b*

Surprisingly, survival to maturity has not been incorporated into calculations of *r*_*max*_ for chondrichthyans (sharks, rays, and chimaeras). As most of these species lack stock-recruitment relationships, survival to maturity at low population sizes has been assumed to be very high and hence set to one because they have high investment per offspring (Dulvy et al., 2014; García et al., 2008; Hutchings et al., 2012). In other words, species with one or hundreds of offspring annually were assumed to have the same survival through the juvenile life stage. However, juvenile survival is likely to vary among chondrichthyans even in the absence of density-dependence as they have a wide variety of reproductive modes (ranging from egg-laying to placental live-bearing) including some of the longest gestation periods in the animal kingdom (Branstetter, 1990). Sensitivity analyses of age-and stage-structured models show that juvenile survival is a key determinant of population growth (λ), especially for species with low *r*_*max*_ (Cortés, 2002; Frisk et al., 2005; Kindsvater et al., 2016).

To correct for the assumption that all juveniles survive to maturity, here we show how the commonly used equation to estimate *r*_*max*_ was derived and then indicate where juvenile survival is accounted for in the model but has been overlooked. We then introduce a simple updated method for estimating *r*_*max*_ that takes into account juvenile survival. Finally, we re-estimate *r*_*max*_ for 94 chondrichthyans using our updated equation and the same life history parameters used previously (see supplementary material in García et al., 2008), compare our updated *r*_*max*_ estimated with previous ones, and discuss which species’ *r*_*max*_ were previously overestimated.

## 2 Methods

### 2.1 Original derivation of *r*_*max*_

The maximum rate of population increase *r*_*max*_ can be derived from the Euler-Lotka equation in discrete time:

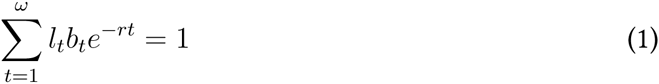
Where *t* is age, *ω* is maximum age, *l_t_* is the proportion of individuals that survive to age *t, b*_*t*_ is fecundity at age *t*, and *r* is the rate of population increase. This rate changes with population density, but we are concerned with the maximum intrinsic rate of population increase *r*_*max*_, which occurs at very low densities in the absense of density dependence. If we assume that after reaching maturity at age *α*_*mat*_ annual fecundity and annual survival are constant (*b* and *p*, respectively), we can estimate the probability of survival to ages *t* >*α*_*mat*_ as survival to maturity *l_αmat_p*^*t–mat*^, where *l*_*αmat*_ is the proportion of individuals surviving to maturity (Myers et al., 1997).

Annual survival of adults is calculated as *p* = *e*^*–M*^ where *M* is the species-specific instantaneous natural mortality rate. This allows for survival to maturity *l*_*αmat*_ and annual fecundity *b* to be removed from the sum and the equation to be rewritten as follows (equation 6 in Myers et al., 1997):

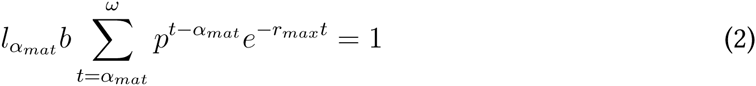

If we solve the summation we obtain the following (see Charnov and Schaffer, 1973; Myers et al., 1997; and Supplementary material for a more detailed derivation)

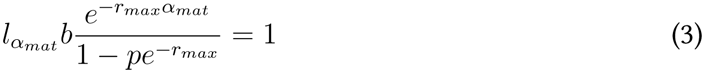
which we can rearrange as

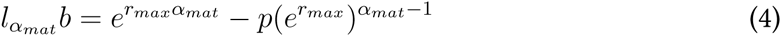

The term outside of the sum *l*_*αmat*_*b* has been equated to the maximum spawners per spawner *ᾶ* thus we can rewrite the equation as

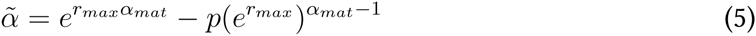

This is the same equation used by Hutchings et al. (2012) to solve for *r*_*max*_ when estimates of *ᾶ* are available, and is mathematically equivalent to the equation used by García et al. (2008) in the case where age of selectivity into the fishery *α*_*sel*_ = 1. Equation 2 shows that survival to maturity is only accounted for in *l*_*αmat*_. Calculations of *ᾶ* for chondrichthyans have ignored *l*_*αmat*_, effectively equating it to 1, assuming *ᾶ = b:*

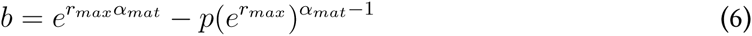

Hence, the previous equation of *r*_*max*_ for chondrichthyans assumed all individuals survived until maturity. This formulation was used for chondrichthyans by García et al. (2008), Hutchings et al. (2012), and Dulvy et al. (2014), and is hereafter referred to as the “previous” equation.

The oversight in the previous formulation of *r*_*max*_ is comparable to an erroneous assumption in fisheries models where steepness—the productivity of the population—is held constant or set to 1 if data from stock-recruitment relationships are not available (reviewed in Mangel et al., 2010). Low-fecundity species such as chondrichthyans are assumed to have extremely high juvenile survival relative to teleost fishes, given that fecundity of sharks and rays is one or two orders of magnitude lower than most teleosts. However, steepness itself is fundamentally a property of early life history traits (Mangel et al., 2010; Myers et al., 1999) and hence should be calculated from demographic data or life history relationships.

Furthermore, it is often assumed that density dependence acts mainly upon juvenile survival. When estimating intrinsic rate of population increase, juvenile mortality is assumed to be lowest at very low population sizes, which may have justified its omission from earlier formulations of the *r*_*max*_ equation (E.L. Charnov, pers. comm.).

### 2.2 Accounting for survival to maturity

We revise the previous method by incorporating an estimate of juvenile survival that depends on age at maturity and species-specific natural mortality. We calculate the proportion of individuals surviving until maturity with the following equation:

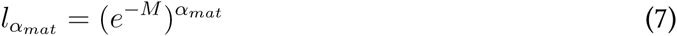

We chose to use a simple estimate of natural mortality *M* based on average lifespan. Assuming that the natural mortality rate of a cohort is exponentially distributed, the average mortality rate is the mean of that distribution, which is equivalent to the reciprocal of average lifespan (Dulvy et al., 2004), such that *M* = 1/*ω*, where *ω* is an estimate of average lifespan, in years (See Supplementary Material). Since cohort data on average lifespan are difficult to obtain, we assume *ω* = (*α*_*max*_ + *α*_*mat*_)/2—the midpoint between age at maturity and maximum age. We do this for three reasons: First, estimates of maximum age are readily available for many chondrichthyan species, and they are applicable to most chondrichthyan populations since they have truncated size class distributions due to prolonged fishing exposure (Law, 2000). Second, chondrichthyans have low fecundity and large offspring, which are much more likely to survive to maturity than species with very high fecundity. This means that the average lifespan and the maximum lifespan are likely much closer together for chondrichthyans than for teleosts. Third, some of the common methods for estimating *M*, e.g., Jensen (1996) or Hewitt and Hoenig (2005), result in unrealistic estimates of *r*_*max*_ for many species (i.e., zero or negative, see Fig. 5 in Supplementary Material) probably due to natural mortality being overestimated for many chondrichthyan species when using estimators based mostly on teleost data. In preliminary analyses we found that when using these teleost-based mortality estimators, we could only obtain plausible estimates of *r*_*max*_ for all species when ignoring juvenile mortality.

One reason for the overestimation may be that the Hewitt and Hoenig (2005) equation coefficients are estimated from data on fish that have extremely low juvenile survival (mostly teleosts). By contrast, our method assumes that 36.8% of offspring reach average lifespan (see explanation and Supplementary Material in Hewitt and Hoenig (2005)). Put simply, for a species with an average lifespan of ten years, 9.5% of the population must die each year for there to be a 37% chance of surviving for ten years. While in teleosts average lifespan is probably less than the age of maturity, for chondrichthyans it is likely greater, which is why we assume it is the mean of age at maturity and maximum observed lifespan. We recalculate *r*_*max*_ for 94 chondrichthyan species examined in García et al. (2008) and Dulvy et al. (2014) using our updated method that combines equations 4 and 7, as well as using the previous method that uses equation 6 and Jensen’s (1996) *M* estimator. Finally, we compare *r*_*max*_ values from previous and updated methods and explore the relationship between life history parameters and discrepancies in *r*_*max*_ values.

## 3 Results and Discussion

Our updated estimates of maximum intrinsic population growth rates (*r*_*max*_) for chondrichthyans are on average 10% lower than previous estimates (Fig. 1). For the most fecund species (*b* > 10 female offspring per year) updated *r*_*max*_ estimates were always 10-20% lower than previous estimates. This means that for species with high fecundity, *r*_*max*_ has been overestimated in the past (see right side of Fig. 2a,b; large circles in Fig. 3). In contrast, for less fecund species (*b* < 5 female offspring per year), discrepancy in *r*_*max*_ between updated and previous estimates varies from 30% lower to 80% higher (small circles in Fig. 3). Two of the most fecund chondrichthyans, the Big Skate *(Raja binoculata)* and the Whale Shark *(Rhincodon typus)*, have lower intrinsic rates of population increase (see Fig. 3) and may be less resilient to fishing than previously thought.

**Figure 1:**
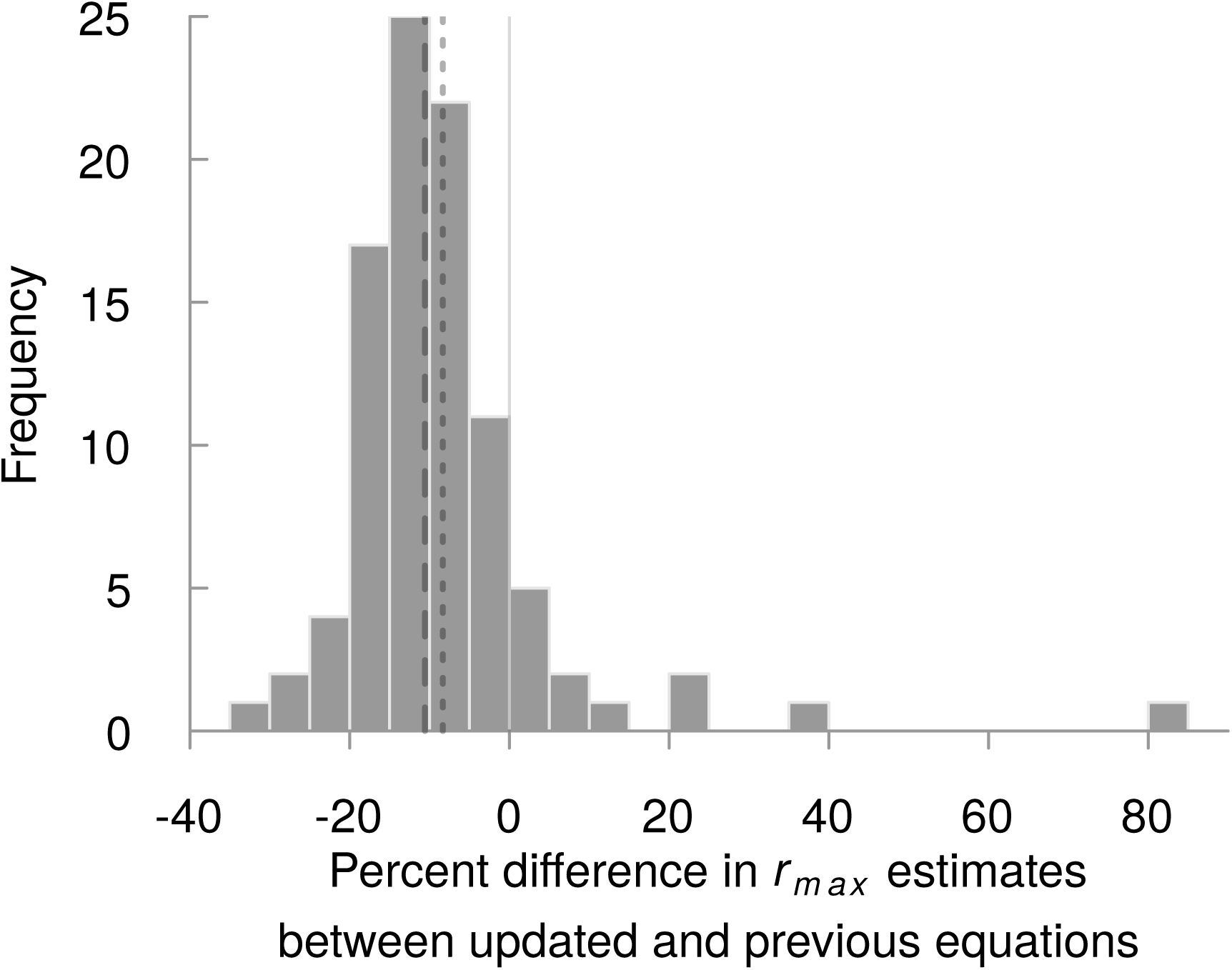
Histogram of percent difference between updated *r*_*max*_ values (this study) and previous ones (from García et al. 2008 and Dulvy et al. 2014). Dashed and dotted lines indicate median and mean values, respectively.

**Figure 2:**
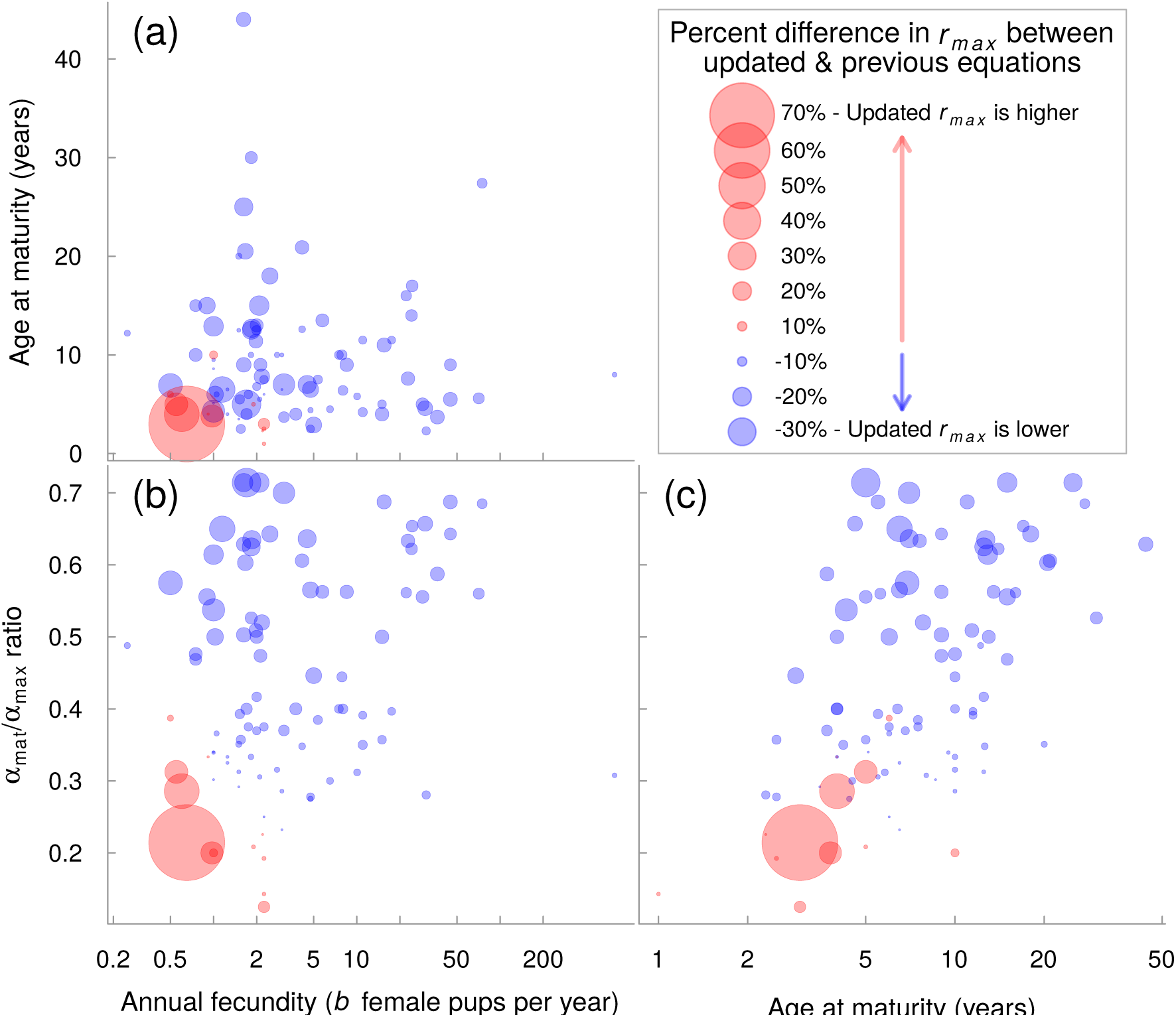
Annual fecundity (*b*, in log-scale) vs (a) age at maturity and (b) the *α*_*mat*_/*α*_*max*_ ratio. (c) Age at maturity vs *α*_*mat*_/*α*_*max*_ ratio. Colour indicates whether the updated model estimates a higher (red) or lower (blue) *r*_*max*_ than the previous formulation, while point size indicates percent difference in *r*_*max*_ estimates between updated and previous models.

**Figure 3:**
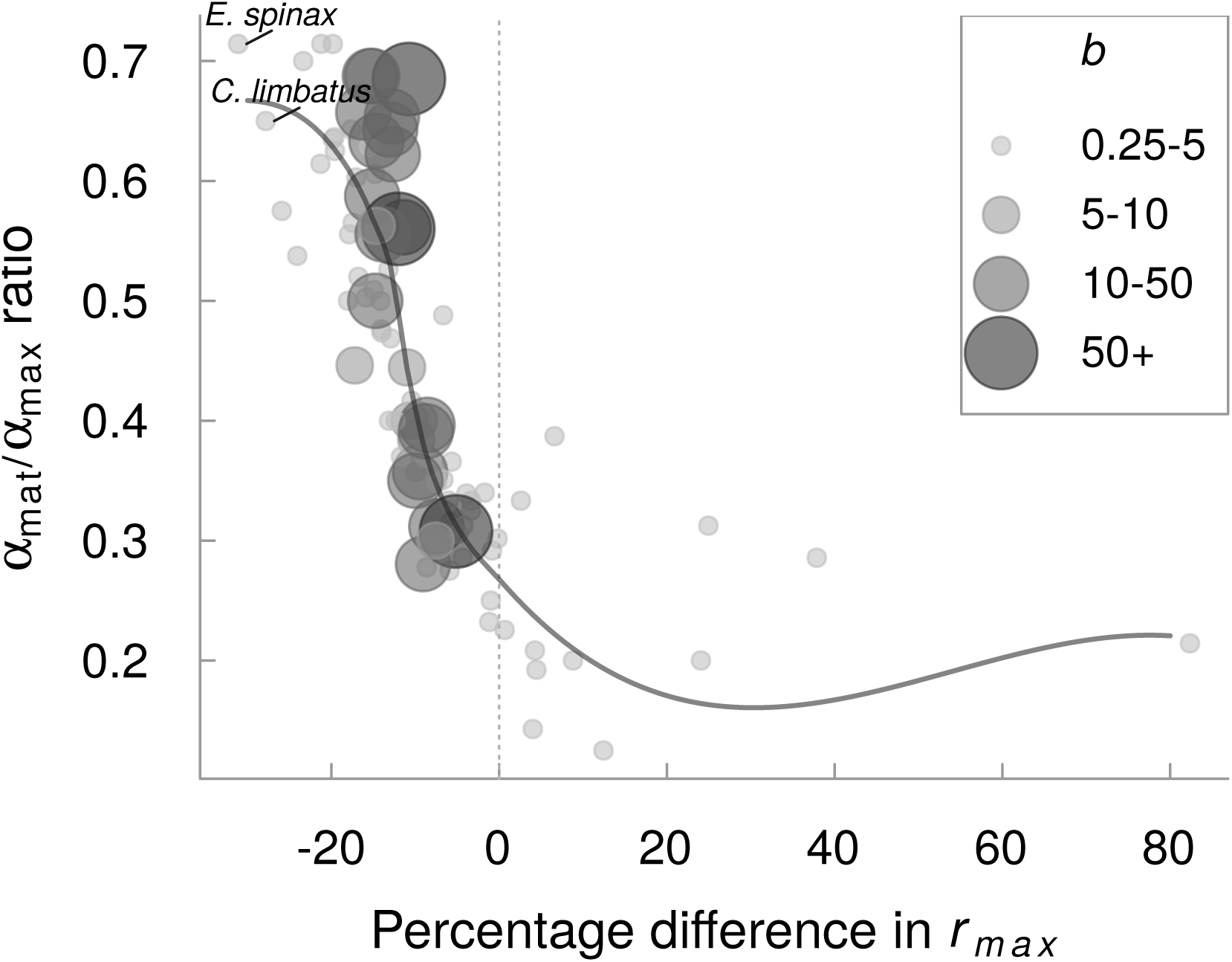
Comparison of percentage difference between updated and traditional *r*_*max*_ and the *α*_*mat*_/*α*_*max*_ ratio across different values of annual reproductive output *b*. Darker grey and larger circles indicate a higher annual reproductive output (b) value. The grey line is the lowess-smoothed curve. Species highlighed are: *E. spinax = Etmopterus spinax, C. limbatus = Carcharhinus limbatus, R. binoculata = Raja binoculata, R. typus = Rhincodon typus*, and *U. lobatus = Urolophus lobatus*.

The greatest positive and negative discrepancies in *r*_*max*_ values (extremes in percent difference) occurred in species with very low annual fecundity and to a lesser extent low age at maturity (see lower left corner of Fig. 2a). The proportional difference between updated *r*_*max*_ and previous estimates were greatest in species with low *r*_*max*_ values. Alternatively, greater fecundity, combined with late maturity “buffer” against variation in estimates of *r*_*max*_ (Fig. 2a,b right side of plots). When age at maturity is low relative to maximum age (*α*_*mat*_/*α*_*max*_ < 0.3), updated *r*_*max*_ estimates were much higher than previous estimates. For example, the updated *r*_*max*_ estimate for the Lobed Stingaree *(Urolophus lobatus)* is 82% higher than its previous *r*_*max*_ estimate, due to its early relative maturation ( *α*_*mat*_/*α*_*max*_ = 0.21, Fig. 3). Conversely, when age at maturity is high relative to maximum age (*α*_*mat*_/*α*_*max*_ > 0.4), updated *r*_*max*_ estimates were lower than previous estimates (Fig. 3). For example, the Velvet Belly Lanternshark *(Etmopterus spinax)* and the Blacktip Shark (*Carcharhinus limbatus*) have relative maturation ratios of 0.71 and 0.65, respectively, and have updated *r*_*max*_ values that are 31% and 28% lower than previously estimated (see Fig. 3). While our study did not explore the relationship between relative maturation (the *α*_*mat*_/*α*_*max*_ ratio) and *r*_*max*_ values among species, a negative relationship between relative maturation and intrinsic rate of population increase has been previously pointed out in sharks (Liu et al., 2015) and skates (Barnett et al., 2013).

Previous work comparing chondrichthyan life histories often overestimated the maximum rate of population increase by not accounting for the species-specific juvenile mortality rate (García et al., 2008; Hutchings et al., 2012). Juvenile survival was overestimated for all species, particularly for highly fecund and late-maturing species, which inflated their estimated maximum intrinsic population growth rates.

Our simple method to estimate survival to maturity requires no extra parameters but assumes that juvenile mortality is equal to adult mortality. This is likely to result in conservative estimates of *M* because juveniles tend to have higher mortality rates than adults (Cushing, 1975). Future work could explore using age-specific mortality estimators to calculate survival to maturity, but we caution that these estimators are mostly based on teleost fishes and require additional data such as on Bertalanffy growth parameters (Chen and Watanabe, 1989) or weight-at-age relationships (Peterson and Wroblewski, 1984).

We found that species with high fecundity all had lower *r*_*max*_ values than previously estimated, hence our method is more effective at representing higher juvenile mortality rates in species with high fecundity. Nonetheless, direct estimates of differential juvenile mortality are still missing from both models, and motivates further research on this topic (Heupel and Simpfendorfer, 2002). Our method undoubtedly ignores nuances related to absolute offspring size and litter size (Smith and Fretwell, 1974), but it is still likely to be an improvement over the previous assumption that all juveniles survive to maturity.

These new insights into the maximum intrinsic rates of increase are relevant for the management of data poor chondrichthyans. We recommend that scientist and managers using chondrichthyan *r*_*max*_ estimates reevaluate them using our updated equation, emphasizing on species whose *r*_*max*_ values have been consistently overestimated in previous studies: highly fecund species, often thought to be more resilient to fishing (Sadovy, 2001), and those that only reproduce during a short span of their total lifetime. To generalize management and conservation implications beyond the species in our study, future work needs to revisit our understanding of life history and ecological correlates of *r*_*max*_. Previous work suggest species in deeper (colder) habitat (García et al., 2008) as well as those with late age at maturity (Hutchings et al., 2012) have lower *r*_*max*_ values. These and other correlates of *r*_*max*_ can now be reevaluated with these updated estimates and used in ecological risk assessments and other forms of management priority setting.

## Acknowledgements

We are grateful to E.L. Charnov and J. Hutchings for discussions on this topic, and to L.K Davidson, P.M. Kyne, J.M. Lawson, and R.W Stein for their comments on the manuscript. This research was funded by the J. Abbott/M. Fretwell Graduate Fellowship in Fisheries Biology (SAP), NSERC Discovery Grants (NKD & JDR), a Canada Research Chair (NKD), and an NSF Postdoctoral Fellowship in Math and Biology (HKK; DBI-1305929).

## Supplementary Material

The Supplementary Material includes a more detailed account on deriving *r*_*max*_ which uses many of the same equations in the main text of the body (here repeated for clarity), details on the conversion of lifetime spawners per spawners to a yearly rate, explanation of why *1/ω* means that 37% of individuals reach average lifespan, and Supplementary Figures.

The raw data used for our analyses are available on figshare at https://dx.doi.org/10.6084/m9.figshare.3207697.v1i.

**Detailed derivation of** *r*_*max*_

The maximum rate of population increase *r*_*max*_ is typically derived from the Euler-Lotka equation in discrete time (Myers et al., 1997):

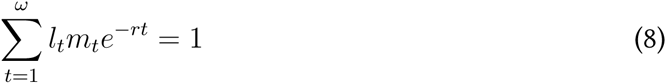

Where *t* is age, *ω* is maximum age, *l*_*t*_ is the yearly survival at age *t, m_t_* is fecundity at age *t*, and *r* is the rate of population increase. This rate changes with population density, but we are concerned with the maximum intrinsic rate of population increase *r*_*max*_, which occurs a very low densities in the absense of density dependence. Assuming that after reaching maturity annual fecundity and annual surival are constant (*b* and *p*, respectively), we can estimate survival to year *t* as survival to maturity *l*_*αmat*_ times yearly adult survival *p* for the years after maturation (Myers et al., 1997):

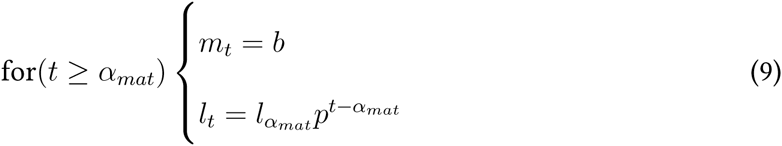
where *α*_*mat*_ is age at maturity, *b* is annual fecundity, and *p* is annual survival of adults and is calculated as *p* = *e*^*–M*^ where *M* is the species-specific instantaneous natural mortality. This allows for survival to maturity *l*_*αmat*_ and annual fecundity *b* to be removed from the sum and the equation to be rewritten as follows (equation 6 in Myers et al., 1997)

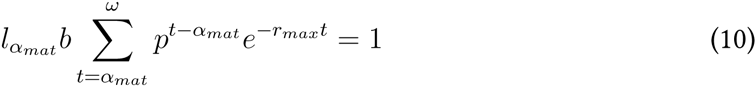

If we assume that *ω* = ∞ we can then solve the geometric series by finding the common ratio. Let *S* be the sum:

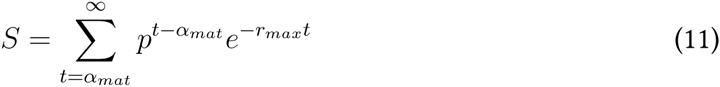

We can break down the summation as:

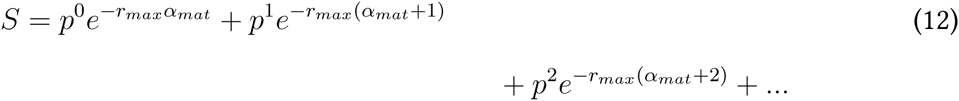
which is equivalent to:

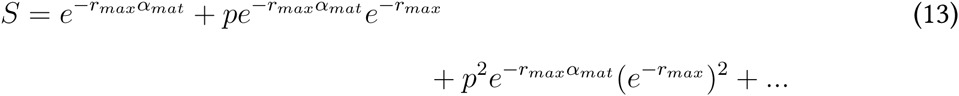

The value that would convert the first item of the sum into the second one, the second item into the third one and so on, is the common ratio, which in this case is *pe*^*-r*_*max*_^. Multiplying everything by *pe*^*-r*_*max*_^ gives us:

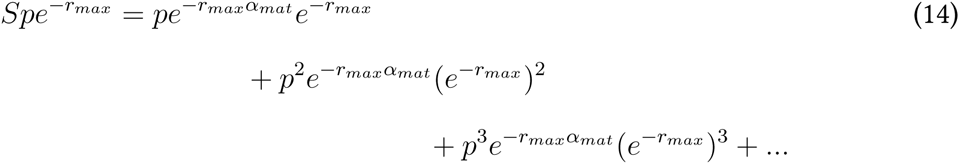

Therefore, the product of *S* and *pe*^*-r*_*max*_^ is equal to *S* minus the first item of the series, *e*^*-r*_*max*_*α*_*mat*_^. We can then subtract this second series (*Spe*^*–r*_*max*_^) from *S:*

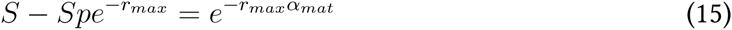

Which allows for estimating *S* as:

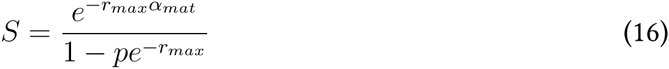

We then replace the summation back in the modified Euler-Lotka equation:

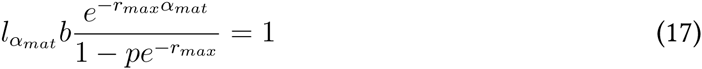

and finally isolate *l*_*αmat*_*b* and rearrange:

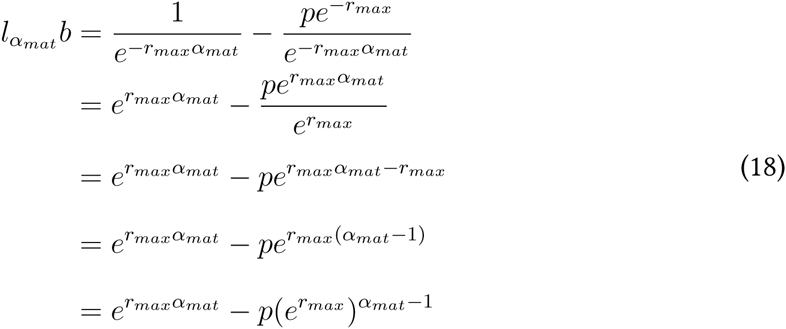

This results in the same equation used by Hutchings et al. (2012), and is mathematically equivalent to the equation used by García et al. (2008) in the case where age of selectivity into the fishery *α*_*sel*_ = 1. Equation 18 shows that survival to maturity is only encapsulated in *l*_*αmat*_ and that its omission effectively assumes that all recruits survive to maturity.

**Understanding why spawners per spawners per year *ᾶ* has been equated with annual fecundity** *b*

All calculations of spawners per spawner are derived from the lifetime spawners per span-wner, 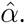 The correct description of *ᾶ* is given in Myers et al. (1997), where it is described as “the number of spawners produced by each spawner per year (after a lag of *α*_*mat*_ years, where *α*_*mat*_ is age at maturity)”. Accounting for that lag is key, as then the lifetime spawners per spawner are divided by the years of sexual maturity, and therefore it is roughly analogous to annual fecundity in females times survival to maturity. The correct way of calculating *a* is by solving

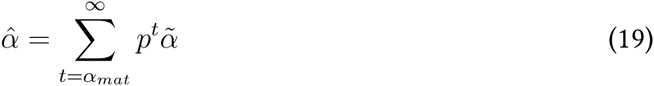

Nonetheless, it has previously been calculated without including the lag of *α*_*mat*_ years, hereafter defined as *ᾶ’*, and has been estimated by solving 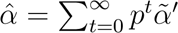, which is the equation used in Myers et al. (1999, 1997) and Goodwin et al. (2006). When using this equation, we are not removing the years before maturity effectively resulting in a metric more akin average yearly spawners per spawner across all age classes. Solving this geometric series without the lag yields:

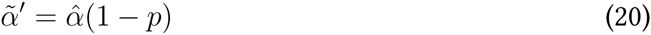

However, as shown in equation 19, we can rewrite the geometric series so that it effectively removes immature age classes. Assuming that after reaching maturity annual surival is constant:

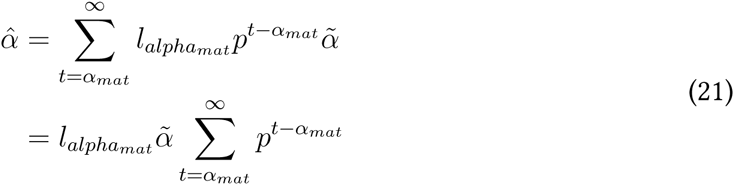

By solving it we obtain the following:

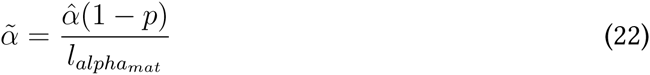
which is analogous to average yearly spawners per spawner across adult age classes, and therefore can be used to estimate *r*_*max*_ instead of *l*_*αmat*_*b*. It also becomes apparent that *ᾶ* = 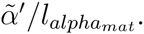. Given that this estimate of *ᾶ* is divided by a proportion, it is larger than the previous estimate; this is expected as lifetime spawners per spawner are partitioned between only by mature age classes (*ᾶ*) instead of all age classes (*ᾶ’*).

**Assumptions of** *M* = 1/*ω*

As already mentioned, we assume that natural mortality rate of a cohort is exponentially distributed, thus the mean of that distribution is the reciprocal of that rate. Estimating instantaneous natural mortality *M* as the reciprocal of average lifespan *ω* is mathematically equivalent to a given percentage of the population reaching *ω*. As previously shown by Hewitt and Hoenig (2005) using their equation as an example, we can calculate that by using *M* as 1/*ω*, we are assuming that, on average, 36.8% of the population reaches average lifespan:

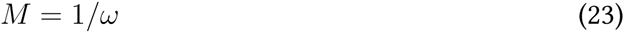

We then rearrange and exponentiate:

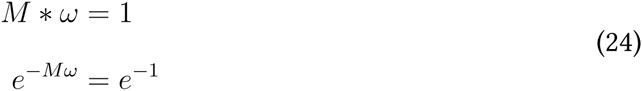

The term *e* ^*–Mω*^ is equivalent to the survival to age *ω* or *lω*. By then calculating the value of *e*^*–1*^ we can see that:

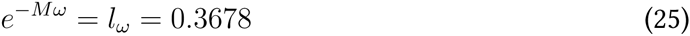

Therefore using our method, the average survival to average lifespan is 36.8%, or roughly one out of three individuals.

**Figure 4:**
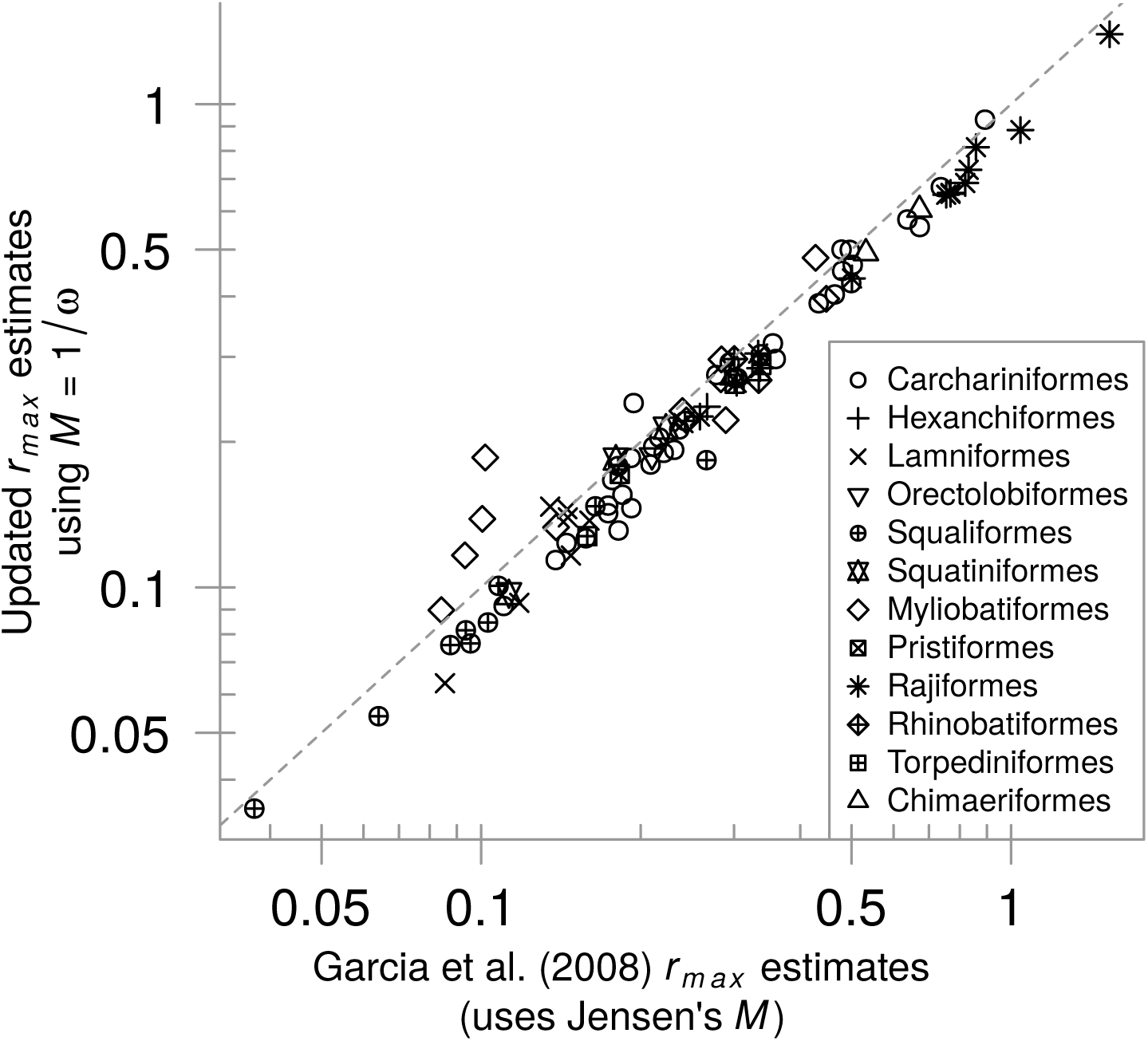
Comparison of previous *r*_*max*_ estimates of the model used in García et al. (2008) (recalculated using the method outlined in their paper) with our updated estimates. Different symbols denote different chondrichthyan orders. Note that the axes are log-transformed.

**Figure 5:**
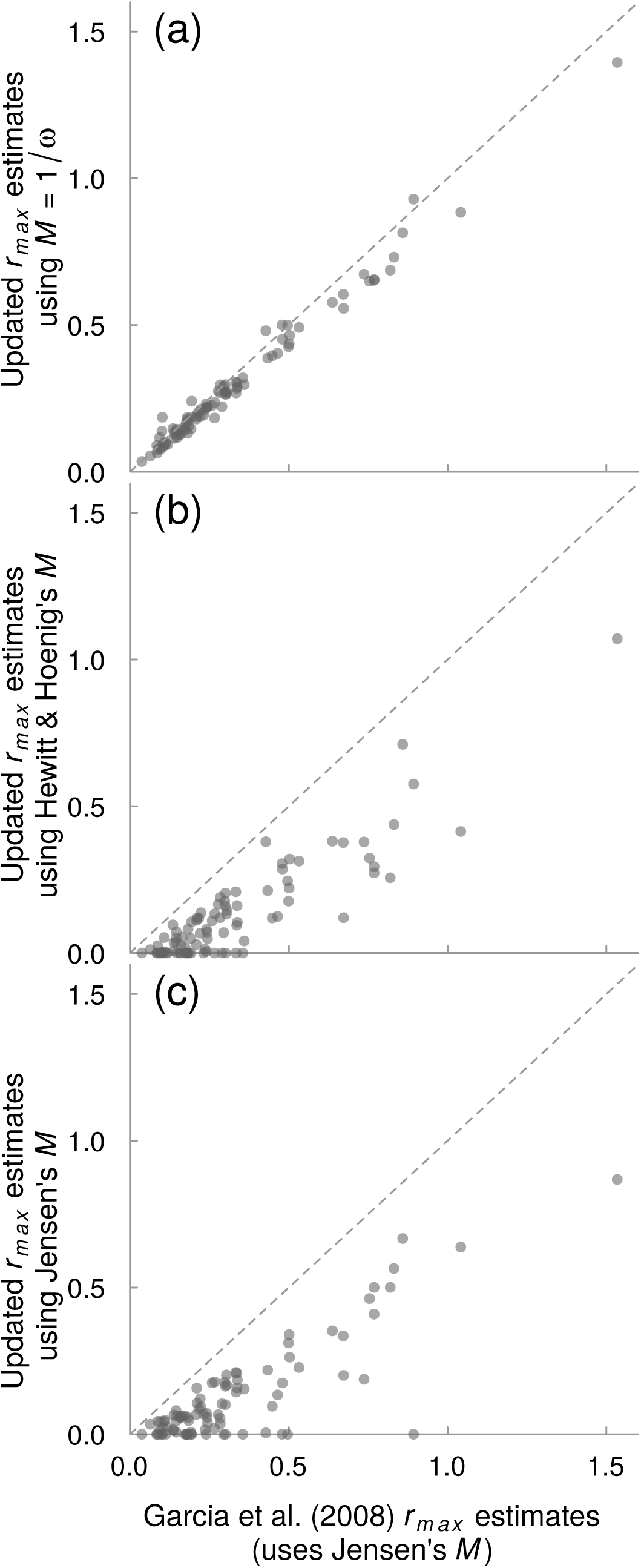
Comparison between updated *r*_*max*_ values with natural mortality estimated from (a) reciprocal of average lifespan, (b) Hewitt and Hoenig (2005), and (c) Jensen (1996). The dashed line represents the 1:1 relationship. Note that only the updated method using the reciprocal of average lifespan (a) shows similar values to the previous *r*_*max*_ estimates, while (b) and (c) often produce *r*_*max*_ estimates equal to zero or negative (both represented here by zeros).

**Figure 6:**
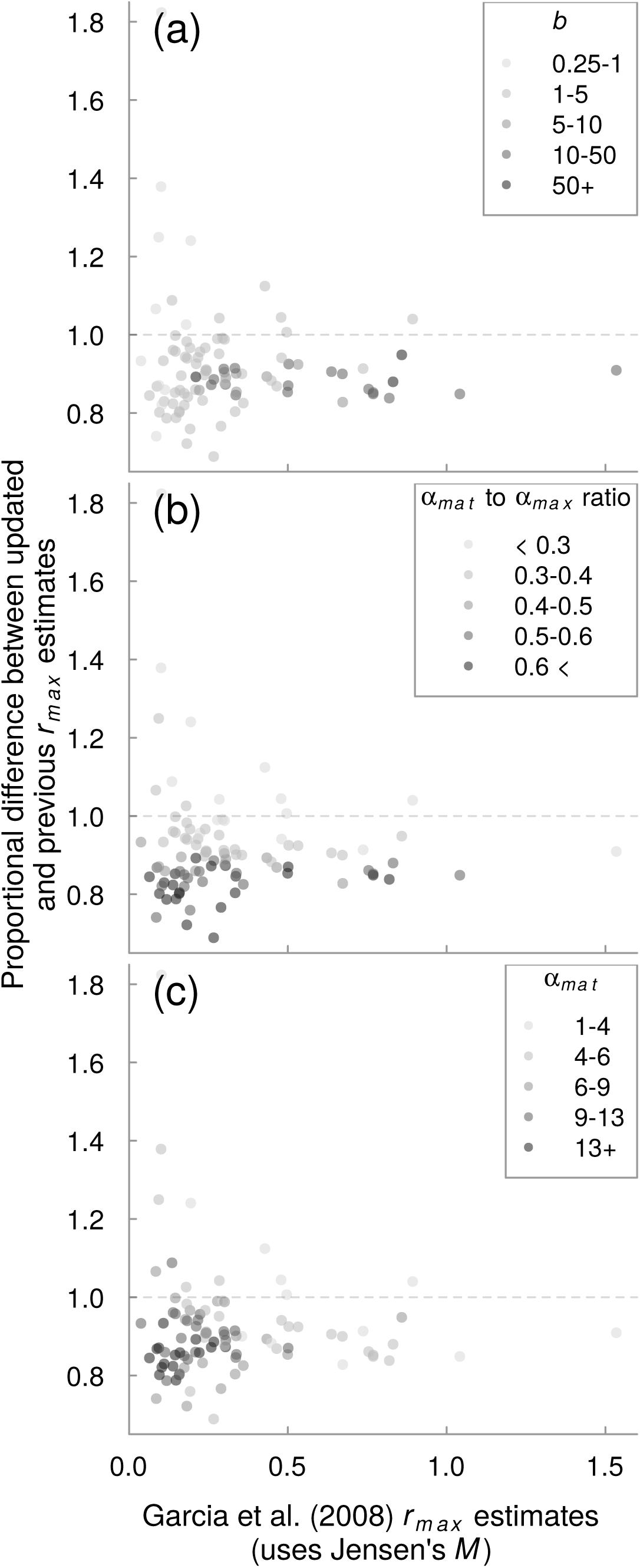
Proportional difference between updated and previous *r*_*max*_ estimates contrasted with (a) annual reproductive output of daughters, (b) *α_mat_*/*α*_*max*_ ratio and (c) age at maturity. The dashed line represents no difference between updated and previous estimates.

## References

Barnett, L.A.K., Winton, M.V., Ainsley, S.M., Cailliet, G.M., and Ebert, D.A. 2013. Comparative Demography of Skates: Life-History Correlates of Productivity and Implications for Management. PLoS ONE 8(5): e65000.

Branstetter, S. 1990. Early life-history implications of selected carcharhinoid and lamnoid sharks of the northwest Atlantic. In Elasmobranchs as living resources: advances in the biology ecology systematics and the status of the fisheries, volume 90, edited by H.L.J. Pratt, S.H. Gruber, and T. Taniuchi, NOAA Technical Report NMFS 90, pp. 17–28.

Charnov, E.L. and Schaffer, W.M. 1973. Life-History Consequences of Natural Selection: Cole’s Result Revisited. The American Naturalist 107(958): 791–793.

Chen, S. and Watanabe, S. 1989. Age dependence of natural mortality coefficient in fish population-dynamics. Nippon Suisan Gakkaishi 55: 205–208.

Cortés, E. 2002. Incorporating uncertainty into demographic modeling: application to shark populations and their conservation. Conservation Biology 16(4): 1048–1062.

Cushing, D.H. 1975. The natural mortality of the plaice. Journal du Conseil 36(2): 150–157. doi:10.1093/icesjms/36.2.150.

Denney, N.H., Jennings, S., and Reynolds, J.D. 2002. Life-history correlates of maximum population growth rates in marine fishes. Proceedings of the Royal Society B 269(1506): 2229–37. doi:10.1098/rspb.2002.2138.

Dulvy, N.K., Ellis, J.R., Goodwin, N.B., Grant, A., Reynolds, J.D., and Jennings, S. 2004. Methods of assessing extinction risk in marine fishes. Fish and Fisheries 5: 255–276.

Dulvy, N.K., Pardo, S.A., Simpfendorfer, C.A., and Carlson, J.K. 2014. Diagnosing the dangerous demography of manta rays using life history theory. PeerJ 2: e400. doi: 10.7717/peerj.400.

Frisk, M.G., Miller, T.J., and Dulvy, N.K. 2005. Life histories and vulnerability to exploitation of elasmobranchs: inferences from elasticity, perturbation and phylogenetic analyses. Journal of Northwest Atlantic Fishery Science 37(October): 27–45. doi:10.2960/J.v35.m514.

García, V.B., Lucifora, L.O., and Myers, R.A. 2008. The importance of habitat and life history to extinction risk in sharks, skates, rays and chimaeras. Proceedings of the Royal Society B 275: 83–89. doi:10.1098/rspb.2007.1295.

Goodwin, N.B., Grant, A., Perry, A.L., Dulvy, N.K., and Reynolds, J.D. 2006. Life history correlates of density-dependent recruitment in marine fishes. Canadian Journal of Fisheries and Aquatic Sciences 63(3): 494–509. doi:10.1139/f05-234.

Heupel, M.R. and Simpfendorfer, C.A. 2002. Estimation of mortality of juvenile blacktip sharks, Carcharhinus limbatus, within a nursery area using telemetry data. Canadian Journal of Fisheries and Aquatic Sciences 59(4): 624–632. doi:10.1139/f02-036.

Hewitt, D.A. and Hoenig, J.M. 2005. Comparison of two approaches for estimating natural mortality based on longevity. Fishery Bulletin 103: 433–437.

Hutchings, J.A., Myers, R.A., García, V.B., Lucifora, L.O., and Kuparinen, A. 2012. Life-history correlates of extinction risk and recovery potential. Ecological Applications 22(4): 1061– 1067. doi:10.1890/11-1313.1.

Jensen, A.L. 1996. Beverton and Holt life history invariants result from optimal trade-off of reproduction and survival. Canadian Journal of Fisheries and Aquatic Sciences 53: 820–822.

Kindsvater, H.K., Mangel, M., Reynolds, J.D., and Dulvy, N.K. 2016. Ten principles from evolutionary ecology essential for effective marine conservation. Ecology and Evolution 6(7): 2125–2138. doi:10.1002/ece3.2012.

Law, R. 2000. Fishing, selection, and phenotypic evolution. ICES Journal of Marine Science: Journal du Conseil 57(3): 659–668. doi:10.1006/jmsc.2000.0731.

Liu, K.M., Chin, C.P., Chen, C.H., and Chang, J.H. 2015. Estimating Finite Rate of Population Increase for Sharks Based on Vital Parameters. PLoS ONE 10(11): e0143008.

Mangel, M., Brodziak, J., and DiNardo, G. 2010. Reproductive ecology and scientific inference of steepness: a fundamental metric of population dynamics and strategic fisheries management. Fish and Fisheries 11(1): 89–104. doi:10.1111/j.1467-2979.2009.00345.x.

Myers, R.A., Bowen, K.G., and Barrowman, N.J. 1999. Maximum reproductive rate offish at low population sizes. Canadian Journal of Fisheries and Aquatic Sciences 56(12): 2404–2419. doi:10.1139/f99-201.

Myers, R.A. and Mertz, G. 1998. The limits of exploitation: A precautionary approach. Ecological Applications 8(1): 165–169.

Myers, R.A., Mertz, G., and Fowlow, P.S. 1997. Maximum population growth rates and recovery times for Atlantic cod, Gadus morhua. Fishery Bulletin 95: 762–772.

Peterson, I. and Wroblewski, J.S. 1984. Mortality rate of fishes in the pelagic ecosystem. Canadian Journal of Fisheries and Aquatic Sciences 41: 1117–1120.

Sadovy, Y. 2001. The threat of fishing to highly fecund fishes. Journal of Fish Biology 59: 90–108. doi:10.1111/j.1095-8649.2001.tb01381.x.

Smith, C.C. and Fretwell, S.D. 1974. The optimal balance between size and number of offspring. The American Naturalist 108(962): 499–506. doi:10.2307/2459681.

